# Expression based biomarkers and models to classify early and late stage samples of Papillary Thyroid Carcinoma

**DOI:** 10.1101/393975

**Authors:** Sherry Bhalla, Harpreet Kaur, Rishemjit Kaur, Suresh Sharma, Gajendra P. S. Raghava

## Abstract

In this study, we describe the key transcripts and machine learning models developed for classifying the early and late stage samples of Papillary Thyroid Cancer (PTC), using transcripts’ expression data from The Cancer Genome Atlas (TCGA). First, we rank all the transcripts on the basis of area under receiver operating characteristic curve, (AUROC) value to discriminate the early and late stage, based on an expression threshold. With the expression of a single transcript DCN, we can classify the stage samples with a 68.5% accuracy and AUROC of 0.66. Then we implemented various combination of multiple gene panels, selected using various gold standard feature selection techniques. The model based on the expression of 36 multiple transcripts (protein coding and non-coding) selected using SVC-L1 achieves the maximum accuracy of 74.51% with AUROC of 0.75 on independent validation dataset with balanced sensitivity and specificity. Further, these signatures also performed well on external microarray data obtained from GEO, predicting nearly 70% (12 samples out of 17 samples) early stage samples correctly. Further, multiclass model, classifying the normal, early and late stage samples achieves the accuracy of 75.43% with AUROC of 0.80 on independent validation dataset. With correlation analysis, we found that transcripts with maximum change in correlation of their expression in both the stages are significantly enriched in neuroactive ligand receptor interaction pathway. We also propose a panel of five protein coding transcripts, which on the basis of their expression, can segregate cancer and normal samples with 97.32% accuracy and AUROC of 0.99 on independent validation dataset. All the models and dataset used in this study are available from the web server CancerTSP (http://webs.iiitd.edu.in/raghava/cancertsp/).

## Introduction

Last few decades witnessed a sharp upsurge in the prevalence of thyroid cancer worldwide making it the fifth most common cancer in women. The exposure to radiation and environmental carcinogens are possible reasons implicated for its rise (Pellegriti et al., 2013). Histopathologically, there are four types of thyroid cancer, stated as Papillary, Follicular, Medullary and Anaplastic. Together, PTC (Papillary Thyroid Carcinoma) and Follicular Thyroid Carcinoma (FTC) are known as Differentiated Thyroid Cancer (DTC) and constitute the majority of thyroid malignancy as well as the most common endocrine malignancy (Hay et al., 2002). According to American Cancer Society, the early stage (stage I and stage II) survival rate of PTC is nearly 100% but that reduces to 55% in stage 4 (American Cancer Society, Cancer Facts & Figures 2017). This statistics already opinions the need for early detection of thyroid cancer.

Fine needle aspiration (FNA) biopsy of the thyroid nodule along with subsequent cytological categorization is a reference method for thyroid cancer diagnosis. It has been reported that the diagnostic precision of FNA has been subjected upon the skill of operator, intrinsic characteristics of nodules and cytology interpretation (Haider et al., 2011). One of its limitations includes that its limited capability to identify follicular lesions (Gharib, 1994). Due to various limitations of FNA cytology, several immunohistochemical markers have been projected and their efficacy in thyroid cancer diagnosis, is being evaluated. Hector battifora mesothelial antigen-1 (HBME-1) (Nga et al., 2008) and Galectin-3 (GAL-3) (Chiu et al., 2010) have shown promising results as diagnostic biomarkers. The severity of thyroid cancer has been shown to correlate with overexpression of EGFR (Sethi et al., 2010). In spite of accumulating knowledge of genetic alterations accompanying the thyroid cancer incidence in the last 20 years (Zolotov, 2016), genomics-based thyroid cancer diagnosis is yet to be realized. The stage wise genome expression analysis can complement the understanding of cancer progression and its correlation to clinical characteristics. The increasing availability of big data has paved the path for deeper understanding of cancer biology in terms of clinical diagnostic and therapeutic capability. One such resource is the TCGA, a public endeavor aimed at establishing a comprehensive catalog of genomic alterations occurring in cancers inferred from large-scale genome sequencing of cancerous tissues accompanied by multidimensional analyses (Tomczak et al., 2015). The big and diverse data generated by TCGA, available to the scientific community, is an incentive to understand the cancer progression and proliferation that would help in devising cancer management solutions. Among the various types and levels of cancer data like genomic mutations, copy number variations, gene fusions, etc., the mRNA expression data is also available for thyroid cancer tissues as well as the associated normal tissues.

The principal paper of PTC from TCGA discussed the genomic landscape of thyroid cancer and identified imperative driver mutations. It also proposed the reclassification of thyroid cancers into molecular subtypes for better understanding of underlying mechanisms (Cancer Genome Atlas Research, 2014). Many other studies have tried to reanalyze this data to study survival, progression and differential expression (Chai et al., 2016;Stokowy et al., 2016;Choi et al., 2017;Yi et al., 2017). Chai et al have shown that the expression of BRAF is associated with high tumor aggressiveness regardless of the BRAF mutation status. So this indicates that the expression and mutation status are both important in determining the prognostic risk factors. Other study illustrates the classification of benign and malignant thyroid tumors on FNA samples with the specificity of 84% (Chudova et al., 2010) on the test dataset of 48 patients. In literature, it has been shown that methylation status of markers like RASSF1, DAPK1 and ESR1 has been significantly associated with thyroid cancer subtypes and early detection of thyroid cancer (Stephen et al., 2015). It has also been revealed that high expression of vitamin D receptor gene (VDR) is associated with classic and tall cell subtype, stage IV and low recurrence-free survival of thyroid cancer (Morris et al., 2010). It has been shown previously that transforming growth factor, cadherin 1, collagen α1, catenin α1, integrin α3, and fibronectin 1 were differentially expressed between benign and malignant nodules of thyroid cancer (Borrelli et al., 2016).

In the present study, we have scrutinized the important transcripts that play an important role in development of cancer from early to late stage using various types of bioinformatics analysis. First, we ranked RNA transcripts based on their discriminatory power to classify early and late stage samples on the basis of an expression threshold. Their gene ontology and pathway analysis is done to ascertain the role of important transcripts in transitioning from early to late stage. Next, multiple transcripts were used to develop models which can segregate early and late stage samples. Further, we have developed a multiclass model to distinguish the normal, early and late stage samples. Moreover, we also performed the correlation analysis to understand the changes in correlation between different transcripts in early and late stage. Next, we have tried to deduce the signature with the minimum number of transcripts that distinguish cancer and normal samples with high accuracy. Additionally, we provide a public domain webserver (CancerTSP) for discriminating the early and late stage along with cancerous or non-cancerous state of the sample using machine learning models.

## Methods

### Datasets

The RNA-Seq data (HTSeq-FPKM, 500 cancer samples and 58 normal) was retrieved from GDC data portal (https://portal.gdc.cancer.gov/). In addition, manifest, metadata, clinical data, biospecimen files were also downloaded from GDC data portal to obtain clinical information using Biospecimen Core Resource (BCR) IDs of patients. For every patient, mRNA expression of 60,483 RNA transcripts were reported in terms of FPKM values. To ascertain the importance of different type of transcripts, we segregated transcripts into subtypes using annotation from genecode version 22; like Protein Coding, LincRNA, snoRNA, snRNA and miRNA etc. transcripts (Supplementary Table S1).

#### Datasets for classification of early vs late stage

Of total 500 cancer samples, there were 281 - stage 1, 52 - stage 2, 112 - stage 3 and 55 - stage 4 samples. In order to develop stage classification method, we divided the samples in early and late stage. Stage 1 and stage 2 samples were designated as early stage samples and stage 3 and stage 4 samples were labelled as late stage samples. Thus, our stage classification dataset contain 333 early and 167 late stage samples. We divided this dataset into training and independent validation dataset where 80% samples were used for training the model and 20% samples for independent validation. Clinical features of the patients are shown in Supplementary Figure S1.

For external validation on the best set of features obtained, we used data from the Gene Expression Omnibus GEO database (Clough and Barrett, 2016) with GEO accession GSE48953. GSE48953 consists of expression profiling of 20 PTC patients using high throughput sequencing (RNAseq) (Smallridge et al., 2014). This dataset consists of 17 early stage patients (stage 1) and 3 late stage patients (stage 3).

#### Other datasets

In addition to stage classification, we also developed model for discriminating cancer patients and normal tissues. This dataset comprises of 500 cancer samples and 58 normal samples. Similarly, we also developed models for predicting normal, early and late stage. The dataset for multiclass classification comprises of 58 normal, 333 early stage and 167 late stage samples. These datasets were further subdivided into training and independent validation dataset, where training dataset contained 80% samples and independent validation dataset contained 20% samples.

### Data processing

The FPKM values were log2 transformed after addition of 1.0 as a constant number to each of the value. After that, we removed features with low variance (less than 0.25) by employing caret package in R, followed by z-score normalization of data. The equations (1) and (2) were used for log transformation and normalization of data, respectively.

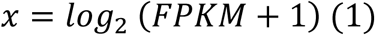

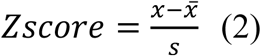

In equation (2), Zscore is the normalized scaled and centered score, x is the log-transformed transcript expression, 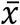 is the mean of transcript’s expression in the training dataset, and s is the standard deviation of a transcript in the training dataset. The log2 transformed independent validation data is z normalized by taking mean and standard deviation of training features.

#### Features filtering using threshold-based models

In this study, we employed a simple expression threshold based approach similar to our previous study (Bhalla et al., 2017) to develop threshold-based models. This method is based on the fact that few transcripts are differentially expressed in early stage as compared to the late stage. In this approach, for every transcript, we designated a threshold, which determines whether a sample is in the early or late stage of the cancer. The threshold is selected by iterating from the minimum to maximum expression of that transcript across all the patients. The threshold which gives maximum AUROC of classification between early and late stage sample is reported. If the mean expression of transcript is greater in early stage than late stage and for a given sample, its log2 FPKM value is higher than the given threshold, then we assign that sample as early otherwise as the late stage sample. Whereas, if the transcript’s average log2 FPKM value is greater in late stage as compared to the early stage and for a given sample, if its log2 FPKM is greater than the threshold, then we assign that sample as the late otherwise as the early stage sample. Using similar method AUROC is calculated for cancer vs. normal samples.

#### Feature selection

To further improve the classification accuracy and develop a multiple gene classification models, we used state-of-the-art techniques to select relevant features. First, we performed feature selection by employing attribute evaluator named, ‘SymmetricalUncertAttributeSetEval’ with search method of ‘FCBFSearch’ of WEKA. The FCBF (Fast Correlation-Based Feature) algorithm employs mainly correlation to identify important discriminating features in high-dimensional datasets in reduced feature space (Yu L, 2003). Secondly, we employed sklearn.feature_selection.F_ANOVA method of feature selection using Scikit-learn package (Pedregosa et al., 2011). This method selects the features by computing F-statistics.

Thirdly, for the model which performed best in comparison to other models, we applied two more feature selection methods. One is wrapper approach for feature selection and other is SVC with L1 penalty (Baraniuk, 2007). In wrapper based approach, human opinion dynamics optimizer (Bhondekar et al., 2011;Kaur et al., 2012;Kaur et al., 2013;Matson et al., 2017) has been used as a search algorithm to search through the space of possible feature subsets with the objective of maximizing MCC on the training set. This is an iterative algorithm in which each candidate solution represents a feature subset. The solution is encoded using 100-bit binary vector where 1 and 0 indicate the presence and absence of a feature in a subset, respectively. The quality of features selected is evaluated using SVM and 10-fold cross validation. The details of algorithm can be found in (Bhondekar et al., 2011;Kaur et al., 2012). The algorithm has been implemented in MATLAB^®^ using LIBSVM and CODO (an open source library hosted on https://github.com/rishemjit/CODO).

### Implementation of machine learning techniques

We have developed machine learning based models using two software; Scikit-learn package and WEKA (Frank et al., 2004). We employed SVM using Scikit-learn and used RBF kernel of SVM at different parameters; g ∈ [10-3 - 10], c ∈ [1-10] using grid search for optimizing the SVM performance. In addition, random forests, SMO, Naïve Bayes and J48 were employed using WEKA software.

### Cross-validation technique

The validation is an indispensable part to evaluate the performance of a prediction method. Ten-fold cross validation technique is exploited to calculate the performance of early vs late stage and cancer vs normal classification models; where, dataset is randomly divided into ten sets, from which nine sets are used as training sets and the left over tenth set as testing dataset. This process is repeated ten times in such a manner that each set is used once as testing dataset.

#### Independent validation

The models developed using 80% data on the best parameters obtained using grid search were used to evaluate independent 20% dataset which was not used for feature selection and training.

### External validation

Further to validate the models on the external validation dataset, first the TCGA data was log2 normalized. Then, the GSE48953 expression data was quantile normalized with reference to TCGA training dataset (target set) using FSQN package (Franks et al., 2018).

### Performance measures

In present study, performance of different models was measured by employing threshold-dependent and threshold-independent parameters. In case of threshold-dependent parameters, sensitivity (Sen), specificity (Spc), overall accuracy (Acc (%)) and Matthews correlation coefficient (MCC) was calculated by using equations (3), (4), (5), and (6) respectively:

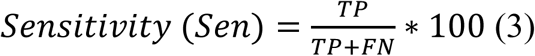

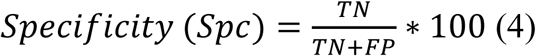

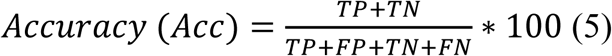

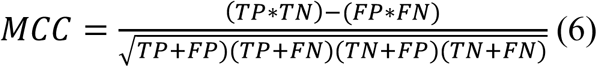

where FP, FN, TP and TN are false positive, false negative predictions, true positive and true negative, respectively.

We also calculated a threshold-independent parameter called AUROC, which is computed from receiver operating characteristic (ROC) plot in this study. The ROC curve is produced by plotting true positive rate against the false positive rate at different thresholds. Lastly, we calculated the area under ROC curve to compute a single parameter from this curve called AUROC in the current study. We used this AUROC value for optimizing and measuring the performance of our models.

In addition, to ascertain the reliability of prediction, we calculated PPV (Positive Predictive Value) and NPV (Negative Predictive Value) at various thresholds using probability score obtained by employing SVM.

#### Multiclass classification

Multiclass classification is implemented using WEKA algorithm using WEKA.classifiers.meta.MultiClassClassifier with random forest as classifier. This method is capable for handling multi-class datasets with 2-class classifiers. This classifier also applies error adjusting output codes for improved accuracy.

#### Functional Enrichment of genes

Enrichment of genes was done using Enrichr tool (Chen et al., 2013;Kuleshov et al., 2016). Only those terms were selected for which adjusted p-value is less than 0.05. Enrichr applies fisher’s exact test along with the adjustment using Bonferroni correction to give adjusted p-values.

## Results

This work has been executed on RNASeq data derived transcript expression from TCGA which consists of 500 PTC and 58 normal tissue samples. Further 500 PTC samples are labelled as stage 1, stage 2, stage 3 and stage 4 samples using clinical information. The stage 1 (281 samples) and stage 2 (52) samples are combined to represent early stage samples and stage 3 (112 samples) and stage 4 (55 samples) samples are pooled to signify late stage samples. The transcript expression is provided in the form of FPKM scores for 60,483 transcripts for all the 500 patients. These transcripts belong to various categories such as protein coding transcripts, long non-coding RNAs, pseudogenes etc. The main objective of this study is to discriminate between early and late stage samples using genome expression data. The overall workflow used in this study is presented in Figure 1 and results are explained in following sections.

**Figure 1:**
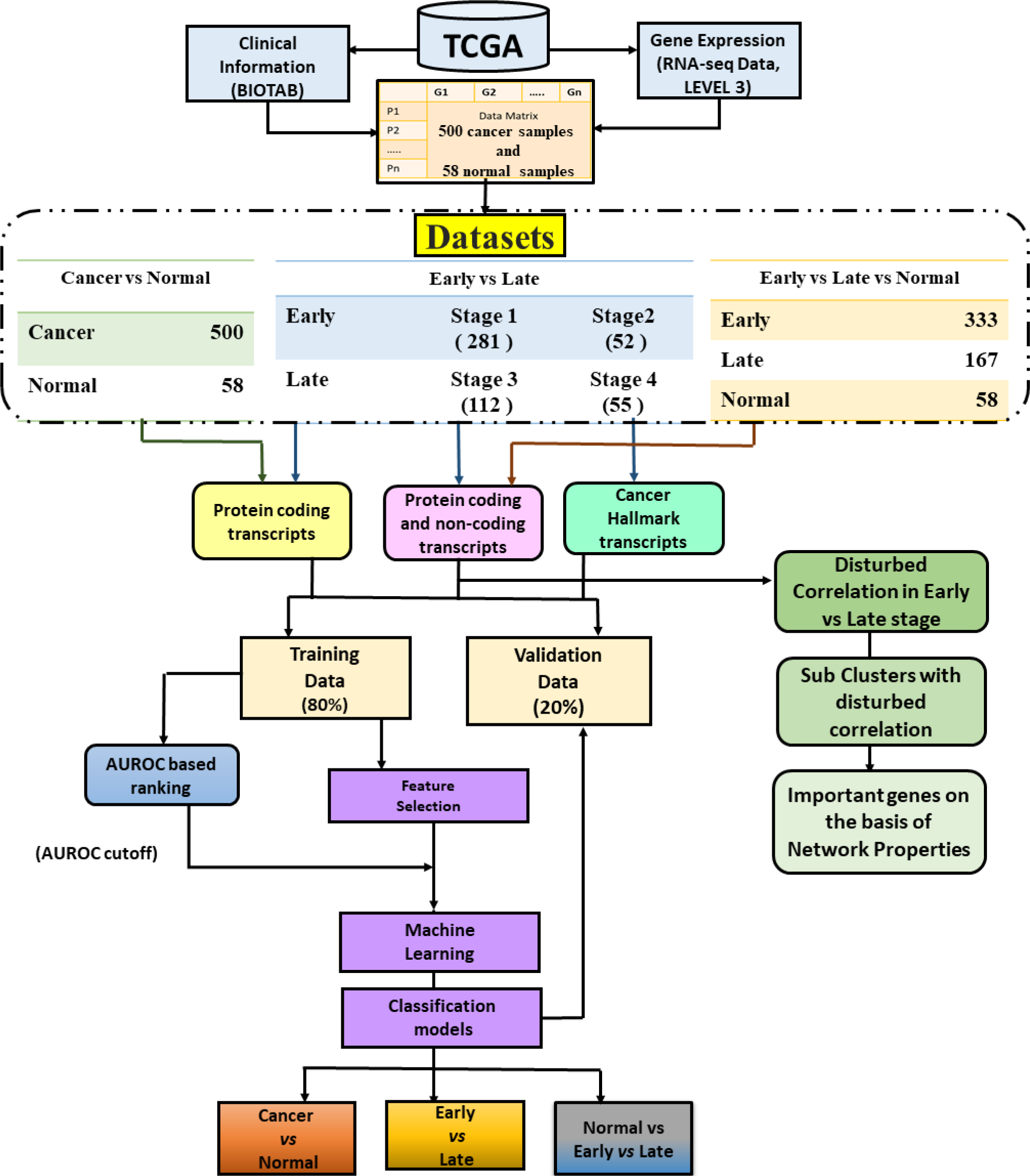
Flowchart representing the overall flow of study, including the types of feature spaces explored for development of machine learning models.

### Single RNA transcript based stage classification

In order to identify classification potential of each RNA transcript, we developed stage classification method using the expression threshold of each RNA transcripts (See Methods). This is a simple threshold based method, where a stage is assigned to a sample if expression of a RNA transcript is more than a given value called threshold; in case the RNA transcript is over expressed in samples of that stage. In order to rank RNA transcripts, we computed the performance of single RNA transcript based models in term of AUROC score. There are 179 transcripts (THCA-EL-AUROC) which have AUROC score greater than equal to 0.60. The THCA-EL-AUROC panel contains key transcripts which can help to discriminate early stage samples from late stage samples and thus can be elucidated as potential biomarkers (Supplementary Figure S2). The DCN protein (overexpressed in late stage) coding transcript shows highest AUROC of 0.66 with accuracy of 68.5%. It is a proteoglycan and its role is well established in discriminating benign and metastasis thyroid and other tumors (Arnaldi et al., 2005;Salomaki et al., 2008). Out of 179 transcripts, 166 are protein coding and 6 are lincRNA and other 7 belong to other classes of non-coding transcripts (Supplementary Table S2).

The 166 protein coding transcripts are significantly represented in 9 oncogenic signatures from MSigDB Database (Liberzon et al., 2015) (Supplementary Table S3) which points out these genes have also been previously implicated in many cancers. The 166 protein coding transcripts are enriched in many pathways of KEGG database such as Focal adhesion pathway (5% genes, adjusted p-value= 4.0e-5), PI3K-Akt signaling pathway (3.5% genes, adjusted p-value=0.001), Proteoglycans in cancer (3% genes, adjusted p-value=0.007). Focal adhesion kinase has been already shown to be overexpressed in thyroid cancers (Kim et al., 2004). There is plethora of literature that shows, PI3K-Akt signaling pathway components are dysregulated in cancers (Fresno Vara et al., 2004;Liu et al., 2009;Matson et al., 2017). In cancer, there is an extensive remodeling of tumor stroma which is related with noticeable variations in proteoglycans expression and structural variability. Proteoglycans mainly contribute to the formation of a matrix for tumor growth affecting tissue organization (Theocharis and Karamanos, 2017). The enriched terms of gene ontology for 166 protein coding transcripts are mainly related to matrix organization and collagen binding (Supplementary Figure S3). Six long non-coding RNAs are found whose AUROC score is greater than equal to 0.6. Thus, literature and enrichment analysis shows that these prioritized genes are involved in various cancer progression related processes and thus can be potential biomarkers of stage classification.

### Stage classification model using multiple RNA transcripts

As shown in above section and in Supplementary Table S2, individual 179 RNA transcripts (THCA-EL-AUROC) have limited ability to classify early and late stage samples with AUROC more than 0.66. Therefore, to develop a model, that can classify stage of samples with high accuracy, we used expression of multiple RNA transcripts. The models based on different machine learning techniques were developed for stage classification using THCA-EL-AUROC (179 RNA transcripts) as features. As shown in Supplementary Table S4, we achieved accuracy of 69.35% with AUROC of 0.72 on training data and accuracy of 67.65% on independent validation data with AUROC of 0.70. The models developed using THCA-EL-AUROC only improved performance marginally from AUROC from 0.66 to 0.70. This means rank-based features are still not sufficient for developing prediction models.

### Protein coding transcripts

As from the previous results, it is evident that protein coding transcripts were major type of transcripts in THCA-EL-AUROC signature, therefore in this analysis we selected 19,814 protein coding transcripts from 60,483 transcripts and applied different feature selection techniques on protein coding transcripts. The FCBF-WEKA (Fast correlation based feature selection method present in WEKA) (Eibe Frank, 2016) based and F_ANOVA (Pedregosa et al., 2011) based feature selection was used to develop models. The SVM model based on 37 features (selected by FCBF-WEKA) is top performer with accuracy of 75.38% and 0.79 AUROC on training data and 69.61% accuracy with AUROC of 0.66 on independent validation (THCA-EL-PC, Table 1) using 37 features obtained using SVM. There was marginal increase in the performance accuracy but the number of features was reduced to reasonable extent as compared to THCA-EL-AUROC. The performance using other algorithms like random forest, Naïve Bayes, SMO and J48 is also shown in Table 1.

**Table 1:** Performance measures of 37-protein coding mRNA feature set (THCA-EL-PC) selected by FCBF-WEKA feature selection method on training and independent validation dataset by using various machine-learning algorithms.

|  | Training |  |  |  |  | Independent validation |  |  |  |  |
| --- | --- | --- | --- | --- | --- | --- | --- | --- | --- | --- |
| Technique | Sens<br>(%) | Spec<br>(%) | Acc<br>(%) | MCC | AUROC | Sens<br>(%) | Spec<br>(%) | Acc<br>(%) | MCC | AUROC |
| SVM | 82.64 | 60.9 | 75.38 | 0.44 | 0.79 | 83.82 | 41.18 | 69.61 | 0.27 | 0.66 |
| Random<br>Forest | 71.70 | 70.68 | 71.36 | 0.40 | 0.77 | 69.12 | 55.88 | 64.71 | 0.24 | 0.63 |
| Naïve<br>bayes | 71.32 | 76.69 | 73.12 | 0.46 | 0.80 | 73.53 | 52.94 | 66.67 | 0.26 | 0.65 |
| SMO | 92.08 | 48.87 | 77.64 | 0.47 | 0.70 | 88.24 | 29.41 | 68.63 | 0.22 | 0.59 |
| j48 | 67.92 | 64.66 | 66.83 | 0.31 | 0.66 | 75.00 | 47.06 | 65.69 | 0.22 | 0.64 |

Sens=Sensitivity; Spec=Specificity; Acc=Accuracy; MCC=Matthews Correlation
Coefficient; AUROC=Area under Receiver operating characteristic curve

We performed interaction analysis in STRING database (Szklarczyk et al., 2015) (Supplementary Figure S4) with THCA-EL-PC transcripts. On adding not more than 10 indirect nodes, we observed three important clusters enriched in different pathways. HIST1H2BJ, the transcript present in our signature forms a cluster and this cluster is enriched in nucleosome cellular component. This cluster has also shown to be related with progression of prostate cancer (Xu et al., 2017) in literature. Another cluster of three genes is enriched in dihydrolipoyl dehydrogenase complex FDR <0.01), out which DBT is present in our original signature. In addition, one more cluster of three genes is a part of checkpoint clamp complex out of which RAD1 is present in original signature and is involved in DNA damage response (Hwang et al., 2015).

In addition, the top 100 features (Supplementary Table S5) were selected using F-ANOVA feature selection method. The accuracy of 71.11% with AUROC of 0.78 is obtained on training data and 69.61% accuracy is obtained using SVM on independent validation data with AUROC of 0.75 (THCA-EL-F-PC, Supplementary Table S6).

### Cancer Hallmark Based Transcripts

Hanahan and Weinberg uncovered the importance of eight biological processes that play vital role in tumor growth and metastatic propagation and called them as cancer hallmark processes (Hanahan and Weinberg, 2011). Thus, genes involved in these processes could also act as key signature markers. Here we have made an effort to ascertain important transcripts from the subset of cancer hallmark genes.

To develop prediction models based on the cancer hallmark genes, we segregated 4,814 cancer hallmark specific genes from 60,483 transcripts and applied feature selection and machine learning algorithms on them. The 15 transcripts selected by FCBF-WEKA (THCA-EL-H) from cancer hallmark genes were used to develop models. The accuracy of 67.84% with AUROC of 0.71 is attained on training data, while the accuracy of 68.63% with AUROC score of 0.73 is obtained on independent validation dataset (Supplementary Table S7). Out of 15 transcripts, two transcripts PROC and NLK (adjusted p-value =0.002) are involved in developmental pathway of Wnt signaling, are shown to be dysregulated in cancer (Ishitani et al., 2003). Other two genes CYSLTR1 and ADRB1 are enriched terms for GPCRs (adjusted p-value=0.04). CYSLTR1 has been shown to be upregulated in colon cancer patients and associated with poor prognosis (Savari et al., 2013). Similar performance is obtained on 50 genes selected using F-ANOVA method (THCA-EL-FH, Supplementary Table S8).

### Protein coding and noncoding transcripts

To ascertain the importance of other non-coding transcripts along with protein coding transcripts, all the 60,483 transcripts were used to select features and develop model on selected features. These 78 transcripts (THCA-EL-All-WEKA) were selected using FCBF-WEKA based feature selection algorithm. The best performance on THCA-EL-All-WEKA panel is obtained using SVM with 78.89% accuracy with 0.86 AUROC on training data and 74.51% accuracy and 0.73 AUROC on independent validation dataset (Supplementary Table S9). Out of 78 selected features, 28 are protein coding transcripts, 12 are long noncoding RNA, 12 are antisense transcripts, 11 are processed pseudogenes and other are different non-coding RNAs (Supplementary Table S10).

The feature selection through F_ANOVA based method with top 100 features (Supplementary Table S11) attained accuracy of 71.86% with AUROC score of 0.72 on training data and accuracy of independent validation data of 61.76% with AUROC score of 0.68 (THCA-EL-All-F, Supplementary Table S12).

Additionally, the top 100 features selected using F_ANOVA were further subjected to the second stage of feature selection. In this stage, a wrapper based approach combining human opinion dynamics optimizer and SVM has been employed (see Methods for details). The number of features were reduced to 27 (Supplementary Table S13, THCA-EL-CODO) and achieved accuracy of 72.25% and AUROC of 0.72 on training set and accuracy of 72% and AUROC of 0.73 on independent validation set (Supplementary Table S14).

As the above three feature selection methods and prediction models gave maximum performance using both protein coding and non-coding transcripts, as compared to protein coding and cancer hallmark protein coding transcripts, we employed another feature selection method which selected features using SVC with L1 penalty (see Methods) which resulted in 36 transcripts. This method gave almost similar accuracy but increased the AUROC from 0.73 to 0.75 (THCA-EL-SVC-L1, Table 2) on independent validation data. This model performed best out of all the models developed to discriminate early and late stage samples in terms of number of features and AUROC on independent validation dataset.

**Table 2:** Performance measures of 36-full feature set (THA-EL-All-SVC) selected by SVC-L1 on training and independent validation dataset by implementing various machine-learning algorithms.

| Technique | Training |  |  |  |  | Independent validation |  |  |  |  |
| --- | --- | --- | --- | --- | --- | --- | --- | --- | --- | --- |
|  | Sens<br>(%) | Spec<br>(%) | Acc<br>(%) | MCC | AUROC | Sens<br>(%) | Spec<br>(%) | Acc<br>(%) | MCC | AUROC |
| SVM | 85.66 | 87.22 | 86.18 | 0.71 | 0.93 | 76.47 | 70.59 | 74.51 | 0.45 | 0.75 |
| Random<br>Forest | 77.36 | 72.93 | 75.88 | 0.49 | 0.8 | 69.12 | 58.82 | 65.69 | 0.27 | 0.7 |
| Naïve<br>bayes | 76.23 | 86.47 | 79.65 | 0.59 | 0.87 | 67.65 | 67.65 | 67.65 | 0.34 | 0.73 |
| SMO | 95.09 | 77.44 | 89.2 | 0.75 | 0.86 | 86.76 | 52.94 | 75.49 | 0.42 | 0.7 |
| j48 | 67.17 | 60.9 | 65.08 | 0.27 | 0.66 | 75 | 50 | 66.67 | 0.25 | 0.62 |

Sens=Sensitivity; Spec=Specificity; Acc=Accuracy; MCC=Matthews Correlation
Coefficient; AUROC=Area under Receiver operating characteristic curve

Further, to ascertain its capabilities, we calculated PPV and NPV on various thresholds of SVM probability score (Table 3). On training data, at the SVM score, greater than 0.9, 161 early stage samples are correctly predicted out of total 170 samples predicted as early stage samples (PPV=94.71%). In case of late stage samples, 60 out of 64 late stage predicted samples are correct (NPV=93.75%). In case of independent validation data, the PPV is 85.71% and the NPV is 66.67% (Table 3). This shows that at SVM score of 0.9, there is high probability of correct positive (early stage) and negative (late stage) prediction. At threshold of 0.7, at which we presented other performance measures in Table 2, the PPV for training data is 93.03% and NPV is 94.51% whereas in case of independent validation the PPV is 83.87% and NPV is 63.64% (Table 3).

**Table 3:** The performance of SVM based model at different threshold in term of probability of correct prediction, developed using 36-full feature set (THA-EL-All-SVC) on training and independent validation dataset.

| Threshold/<br>Cut-offs | Prediction of Early Stage |  |  | Prediction of Late Stage |  |  |
| --- | --- | --- | --- | --- | --- | --- |
|  | Total<br>Predictions | Correct<br>Prediction | PPV | Total<br>Predictions | Correct<br>Prediction | NPV |
| Performance of Training Dataset |  |  |  |  |  |  |
| 1.00 | 16 | 16 | 100.00 | 6 | 6 | 100.00 |
| 0.95 | 136 | 131 | 96.32 | 42 | 40 | 95.24 |
| 0.90 | 170 | 161 | 94.71 | 64 | 60 | 93.75 |
| 0.85 | 199 | 188 | 94.47 | 71 | 67 | 94.37 |
| 0.80 | 219 | 207 | 94.52 | 80 | 76 | 95.00 |
| 0.75 | 233 | 221 | 94.85 | 86 | 81 | 94.19 |
| 0.70 | 244 | 227 | 93.03 | 91 | 86 | 94.51 |
| 0.65 | 254 | 235 | 92.52 | 98 | 90 | 91.84 |
| 0.60 | 261 | 240 | 91.95 | 106 | 97 | 91.51 |
| Performance of Independent validation Dataset |  |  |  |  |  |  |
| 1.00 | 0 | 0 | 0.00 | 0 | 0 | 0.00 |
| 0.95 | 28 | 25 | 89.29 | 9 | 7 | 77.78 |
| 0.90 | 42 | 36 | 85.71 | 12 | 8 | 66.67 |
| 0.85 | 47 | 39 | 82.98 | 15 | 9 | 60.00 |
| 0.80 | 54 | 45 | 83.33 | 16 | 10 | 62.50 |
| 0.75 | 58 | 48 | 82.76 | 19 | 12 | 63.16 |
| 0.70 | 62 | 52 | 83.87 | 22 | 14 | 63.64 |
| 0.65 | 65 | 53 | 81.54 | 22 | 14 | 63.64 |
| 0.60 | 70 | 56 | 80.00 | 25 | 16 | 64.00 |

PPV=positive predictive value; NPV=Negative predictive value

We also applied various other state-of-the-art machine learning methods on THCA-EL-SVC-L1, but SVM performed best out of all (Supplementary Table S15). One of the advantage of this method is that it resulted a more balanced sensitivity and specificity along with higher AUROC on less number of features (36 features, THCA-EL-SVC-L1) as compared to 78 features (THCA-EL-All-WEKA) selected by WEKA. These 36 transcripts consists of 17 protein coding genes, 6 long non-coding RNAs and rest other types of non-coding RNA transcripts (Supplementary Table S16). The TERT gene in this signature has been an important oncogene in case of PTC (Liu and Xing, 2016). Overexpression of TERT induced by MAP pathway has shown to aggravate tumor development (Liu et al., 2018).

This model is so far the paramount model in our analysis for binary classification for early and late stage samples. This also points out that both protein coding and non-coding transcripts play an important role in tumor development.

In order to validate these feature on external dataset, we used RNA-Seq data, GSE48953 from GEO database of 20 patients. This dataset contained only 18 features out of 36 features from signature. So, we developed model on quantile normalized 18 features and tested on external validation dataset after applying quantile normalization to it in reference to training data. This model predicted 70.6% (12 samples out of 17 samples) early samples and 100% (3 samples) late samples correctly. This further strengthens and validate our signature on dataset addition to TCGA.

### Multiclass Classification

One of the limitation of binary classification is that it would force even normal samples into either early or late stage samples. Therefore we implemented multiclass classification by taking into account the normal samples available in TCGA for thyroid cancer. We used FCBF-WEKA to select 211 transcripts (THCA-NEL-M, Supplementary Table S17) from 60,483 transcripts and tried to classify normal, early and late stage samples. This model is able to classify 77.02% samples correctly with weighted AUROC of 0.84 in training data and achieved 75.43% accuracy in independent validation data with AUROC of 0.80.

At 0.6 threshold, where coverage is maximum, 44 normal samples out of 45 predicted normal samples are correct. At same threshold, 227 early stage samples are correctly predicted out of 285 total predicted early stage predicted samples. At the same threshold, all the 17 late stage predicted samples are correct (Table 4).

**Table 4:** Confusion matrix for multiclass classification at different thresholds, for each threshold we predict samples of each class, identify correctly predicted in same class and wrongly predicted samples of other classes.

| Threshold /<br>Cut-offs | Prediction for Normal samples |  |  |  | Prediction for Early stage |  |  |  | Prediction for Late stage |  |  |  |
| --- | --- | --- | --- | --- | --- | --- | --- | --- | --- | --- | --- | --- |
|  | Total<br>Predicted | Correctly<br>Predicted | Early | Late | Total<br>Predicted | Correctly<br>Predicted | Normal | Late | Total<br>Predicted | Correctly<br>Predicted | Normal | Late |
| Training dataset |  |  |  |  |  |  |  |  |  |  |  |  |
| 0.90 | 4 | 4 | 0 | 0 | 0 | 0 | 0 | 0 | 0 | 0 | 0.00 | 0 |
| 0.85 | 9 | 9 | 0 | 0 | 3 | 3 | 0 | 0 | 0 | 0 | 0.00 | 0 |
| 0.80 | 19 | 19 | 0 | 0 | 20 | 18 | 0 | 2 | 0 | 0 | 0.00 | 0 |
| 0.75 | 29 | 29 | 0 | 0 | 78 | 65 | 0 | 13 | 1 | 1 | 0.00 | 0 |
| 0.70 | 37 | 37 | 0 | 0 | 154 | 132 | 0 | 22 | 2 | 2 | 0.00 | 0 |
| 0.65 | 42 | 42 | 0 | 0 | 242 | 198 | 1 | 43 | 7 | 7 | 0.00 | 0 |
| 0.60 | 45 | 44 | 1 | 0 | 285 | 227 | 1 | 57 | 17 | 17 | 0.00 | 0 |
| Independent validation dataset |  |  |  |  |  |  |  |  |  |  |  |  |
| 0.90 | 0 | 0 | 0 | 0 | 0 | 0 | 0 | 0 | 0 | 0 | 0.00 | 0 |
| 0.85 | 1 | 1 | 0 | 0 | 0 | 0 | 0 | 0 | 0 | 0 | 0.00 | 0 |
| 0.80 | 5 | 5 | 0 | 0 | 4 | 3 | 0 | 1 | 0 | 0 | 0.00 | 0 |
| 0.75 | 5 | 5 | 0 | 0 | 18 | 15 | 0 | 3 | 0 | 0 | 0.00 | 0 |
| 0.70 | 6 | 6 | 0 | 0 | 43 | 35 | 1 | 7 | 0 | 0 | 0.00 | 0 |
| 0.65 | 6 | 6 | 0 | 0 | 58 | 46 | 1 | 11 | 0 | 0 | 0.00 | 0 |
| 0.60 | 9 | 9 | 0 | 0 | 74 | 58 | 1 | 15 | 4 | 3 | 0.00 | 1 |

An interesting observation was seen when these 211 transcripts were analyzed in STRING. As we added 5 indirect iterations to the network, many of the protein coding transcripts in our signature were interacting with PCNA (Supplementary Figure S5). PCNA has been associated with fatal outcomes (Basolo et al., 1993). The genes in our signature directly interacting with PCNA points out to their importance in acting as stage specific biomarkers.

### Correlation Disturbance in Early and late stages of cancer

To further elucidate the expression of various transcripts in early versus late stage, the Pearson correlation coefficient of each transcript was calculated with other 60,483 transcripts in early and late samples separately. The correlations were compared and only significant correlations were taken into consideration for further analysis. The pairs which had statistical significant correlation difference of 0.70 (adjusted p-value < 0.05) (Supplementary Table S18) were segregated for subsequent analysis based on the assumption that this large change or disturbance in correlation is due to diseased condition. These were the gene pairs whose correlation has been disturbed drastically in early stage as compared to late stage and vice versa. From this analysis, we obtained two types of transcripts i.e. one which had low correlation in early stage patients but high correlation in late stage patients (L_pairs) and others had high correlation in early stage patients and low in late stage patients (E_pairs).

Furthermore, second filter of correlation is also applied on E_pairs and L_pairs. Here, only those E_pairs are taken for further analysis which have correlation value greater than 0.6. There were total 453 pairs which had correlation of 0.60 or higher in early samples and their correlation difference between early and late samples is at least 0.7. Further, gene enrichment analysis indicates that these transcripts are mostly enriched in neuroactive ligand receptor interaction (adjusted p-value=0.01) and folate synthesis pathway (adjusted p-value=0.01).

In addition, there are 778 L_pairs which had correlation greater than 0.60 in late stage and had a shift of correlation at least 0.70 in comparison to early stage. Surprisingly the transcripts present in L_pairs were also enriched in neuroactive ligand receptor interaction (adjusted p-value=0.00002) like E_pairs. Moreover, these genes were also enriched in cellular components like acetylcholine-gated channel complex (adjusted p-value=0.002), ionotropic glutamate receptor complex and G-protein coupled receptor complex. Genome-wide association study has revealed that nicotine acetylcholine receptor (nAChR) cluster was related with lung cancer risk (Niu and Lu, 2014). Glutamate receptors were found to be differentially expressed in case of many cancers like brain tumors (de Groot et al., 2008) and pancreatic cancer (Herner et al., 2011).

Subsequently, the genes of both the categories were combined and network analysis was performed using Cytoscape (Shannon et al., 2003) to understand and elucidate important genes by converting these gene pairs as network. The nodes are the transcripts and their correlation in early stage is taken as edges. As the network is quite complex and has many unconnected components, thus we created subnetworks for better understanding of this network. Accordingly, we analyzed the top subnetwork with 261 nodes of this network on the basis of two main important properties i.e. degree and Betweenness centrality. The Betweenness centrality measures the extent to which a vertex lies on paths between other vertices. Vertices with high betweenness may have extensive effect within a network as such vertices control information passing between others. Their disruption can result in disruption of communication among the network (Pavlopoulos et al., 2011).

In the most populated subnetwork of 261 transcripts, the top five genes having high degree are PRKCG, MNX1, PRSS33, PLK5 and TRH (Figure 2). The importance of these genes was also indicated by the results that shown that these were the genes which have disturbed correlation with maximum number of other genes. The protein kinase C (PKC), a receptor for the tumor-promoting phorbol esters has been hugely studied in the perspective of cancer (Antal et al., 2015). The PRKCG kinase has been described to be involved in cancer cell proliferation and invasion (Korc, 2010). It has also been shown that PRKCG gene involved in tumorogenesis in many cancers (Mazzoni et al., 2003;Parsons and Adams, 2008). The role of PGR has been shown to have role in development of breast cancer (Mohammed et al., 2015). MNX1 as a novel oncogene has shown to be upregulated to a relatively greater degree in prostate cancer (Zhang et al., 2016), acute myeloid leukemia (von Bergh et al., 2006) and hepatocellular carcinoma (Wilkens et al., 2011). This gene also has high Betweenness centrality which points out to be important component of this network. The other genes with high Betweenness centrality are FAM135B and FOXN4. Latter is a member of transcriptional regulators and has been shown to regulate various important processes and has already been shown to promote tumorigenesis (Myatt and Lam, 2007). FAM135B has been implicated in promoting malignancy of ESCC (Eoesophageal squamous cell carcinoma) cells (Song et al., 2014). Thus, this analysis highlights some important genes which are not reflected in expression analysis.

**Figure 2:**
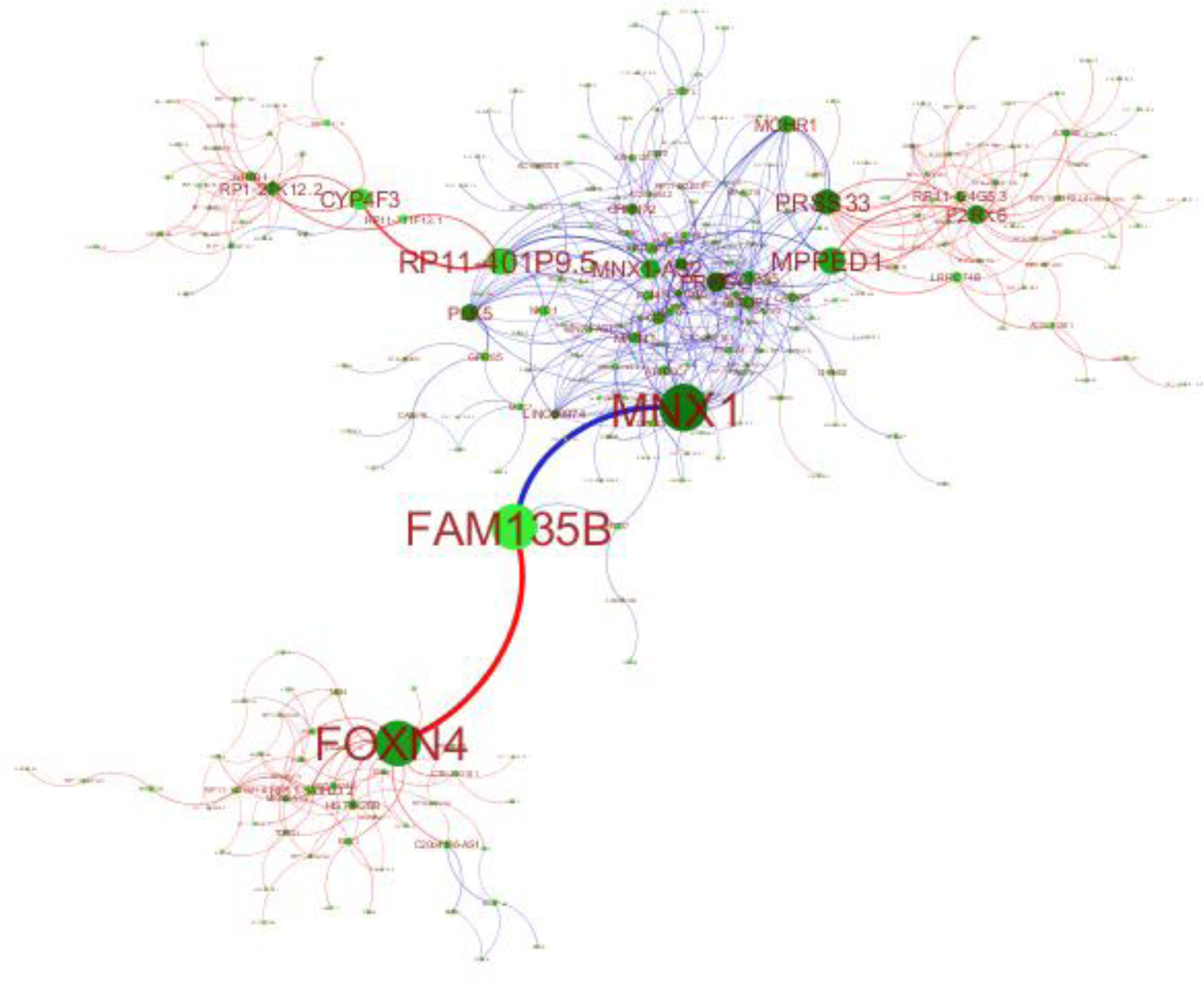
Sub Network of genes which have high shift in correlation (>0.7). Here circle represents nodes and connecting edges represents expression correlation in early stage patients. Dark green color indicates high degree and bigger size of node indicates high betweenness centrality. The blue colored edge indicate correlation less than 0.6 in early stage patients but correlation between these genes is at least shifted by 0.7 in late stage and red colored edges indicate correlation greater than 0.6 in early stage and a shift of at least 0.7 in late stage.

### Discrimination of cancer vs normal samples

#### Single gene based models

After identification of genes/RNA transcripts that can classify early and late stage of PTC, our next goal is identify important RNA transcripts that can distinguish PTC samples from normal tissue samples. Hence, threshold based analysis is done to ascertain features which can discriminate cancer samples from normal. Out of 60,483 features, those features were removed which have variance of less than 0.02, thus number of transcripts reduced to 24,334. Among 24,334 transcripts, there are 8,180 genes which have AUROC of 0.6 or greater. To understand most important genes, we selected only those transcripts which have AUROC of 0.85 or greater (426 transcripts, Supplementary Table S19). The overlapping sense transcript RP11-363E7.4 and protein coding transcript FAM84A shows AUROC of differentiation of 0.96 and 0.95 respectively for classifying cancer samples as compared to normal samples. For 386 protein coding transcripts out of 426 transcripts, pathway, Gene Ontology analysis summary is shown in Supplementary Figure S6. These 386 transcripts contain 11 genes which are involved in axon guidance pathway from KEGG. These genes have been shown to play important role in tumorigenesis (Chedotal, 2007). Enrichment analysis shows that eight genes from 386 signatures have been found in lung-cancer specific 86 genes defined in KEGG. The genes are enriched in biological processes which are regulating expression of non-coding RNAs. The enriched cellular components terms are mostly related to interleukin receptor complexes, T-cell receptor complexes and plasma membrane components. Serum IL-2 has been shown to discriminate patients with active thyroid cancer from the healthy with a sensitivity of 98% and specificity of 58%, (adjusted p-value = 0.0007) (Martins et al., 2018). Other than protein coding genes there are 17 long noncoding RNAs which have AUROC greater than 0.85. Among 17 lincRNAs, PTCSC3, has been shown to be highly thyroid-specific and is downregulated in thyroid tumor tissues and thyroid cell lines (Fan et al., 2013).

LINC00936 and RP11-774O3.3 have been shown to be downregulate in lung cancer (Yu et al., 2015). LINC00205 has been shown to be associated with reoccurrence of hepatocellular carcinoma (Cui et al., 2017). It points out the literature validation of the important signatures found out during this analysis.

### Protein coding RNA transcript based signatures

Our next goal is to develop prediction model based on the least number of protein coding RNA features to classify cancer and normal samples. We selected top five features RELN, RASSF9, PLA2R1, MMRN1 and RPS6KA5 using F-ANOVA (THCA-CN-F). We have developed prediction models for 5 transcripts set using various machine-learning algorithms and they performed reasonably well with SVM with accuracy 98.21%, MCC 0.90 and AUROC 0.97 on training dataset and accuracy 97.32%, MCC 0.88 and AUROC 0.99 on independent validation dataset (Table 5).

**Table 5:**
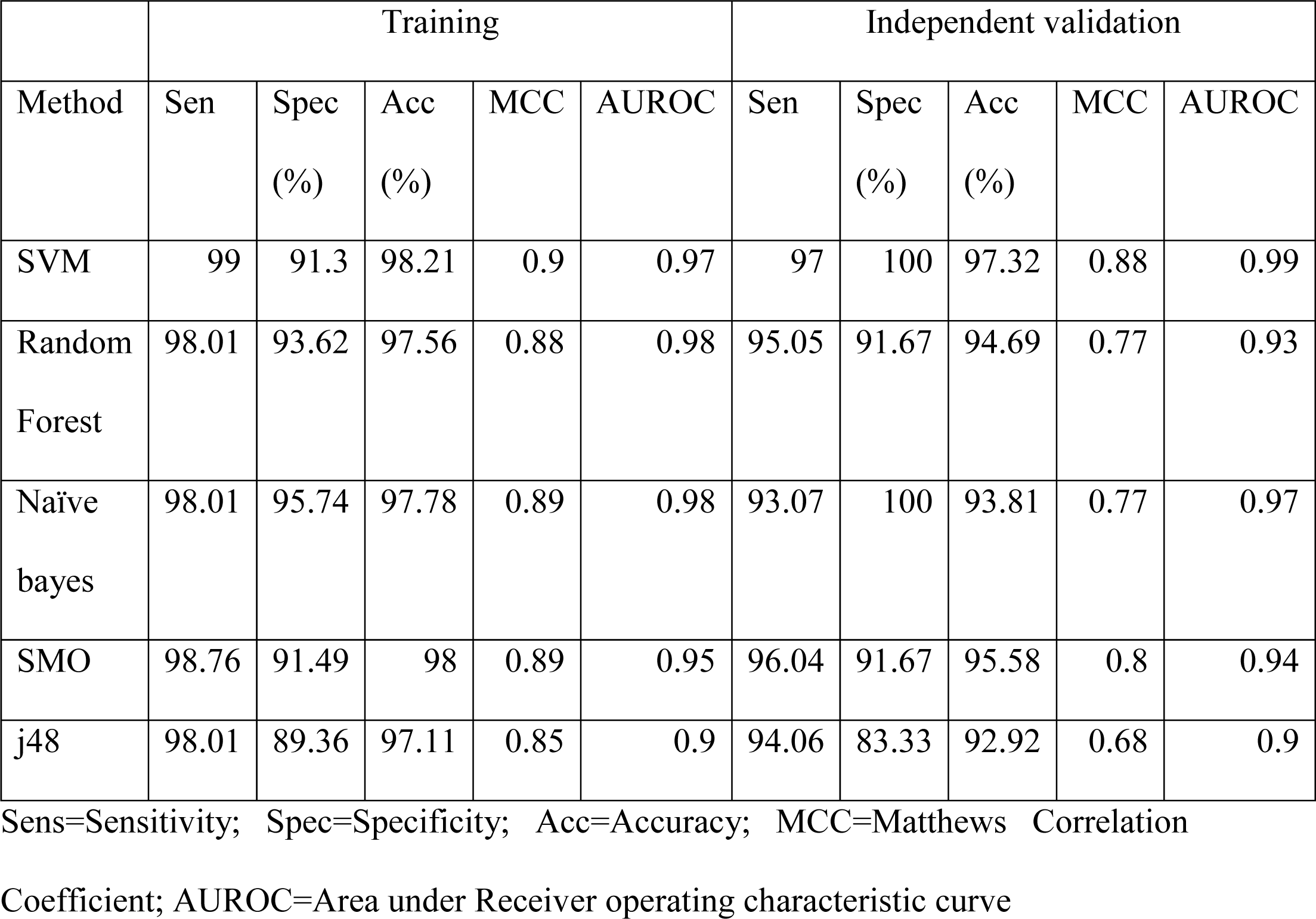
Performance measures of 5-protein coding transcripts (THCA-CN-F) feature set selected by F-ANOVA feature selection method for discriminating cancer and normal patients on training and independent validation dataset.

|  | Training |  |  |  |  | Independent validation |  |  |  |  |
| --- | --- | --- | --- | --- | --- | --- | --- | --- | --- | --- |
| Method | Sen | Spec<br>(%) | Acc<br>(%) | MCC | AUROC | Sen | Spec<br>(%) | Acc<br>(%) | MCC | AUROC |
| SVM | 99 | 91.3 | 98.21 | 0.9 | 0.97 | 97 | 100 | 97.32 | 0.88 | 0.99 |
| Random<br>Forest | 98.01 | 93.62 | 97.56 | 0.88 | 0.98 | 95.05 | 91.67 | 94.69 | 0.77 | 0.93 |
| Naïve<br>bayes | 98.01 | 95.74 | 97.78 | 0.89 | 0.98 | 93.07 | 100 | 93.81 | 0.77 | 0.97 |
| SMO | 98.76 | 91.49 | 98 | 0.89 | 0.95 | 96.04 | 91.67 | 95.58 | 0.8 | 0.94 |
| j48 | 98.01 | 89.36 | 97.11 | 0.85 | 0.9 | 94.06 | 83.33 | 92.92 | 0.68 | 0.9 |

Sens=Sensitivity; Spec=Specificity; Acc=Accuracy; MCC=Matthews Correlation Coefficient; AUROC=Area under Receiver operating characteristic curve

Towards their biological significance, gene enrichment analysis revealed that RELN and RPS6KA5 have been associated with activation of cyclic AMP (cAMP) response element binding protein (CREB) transcription factor (adjusted p-value < 0.01), which is responsible for tumor initiation, progression and metastasis (Cui et al., 2017). RELN is an extracellular glycoprotein that plays a vital role in neuronal migration and has been shown to down regulated in many cancers (Dohi et al., 2010;Okamura et al., 2011). We also selected features using WEKA and achieved similar performance as of 5 features based models (Data not shown).

### Web Server Implementation

We established a web server, CancerTSP (Thyroid cancer stage prediction), that implements models established in the present study for investigation and estimation of cancer stage from the transcripts’ expression data. CancerTSP consists of three key modules; first for prediction whether sample is cancerous or normal, second is for predicting whether it is in early or late stage cancer, and third is for the analysis of transcripts’ expression data.

Prediction module allows the users to predict whether the sample is cancerous or not and also predicts the stage of the cancer using FPKM values. The user needs to provide transcript expression (FPKM values) of potential biomarker genes for every patient. The number of patients corresponds to the number of columns in a file. The output includes a list for patient and corresponding predicting stage of cancer (early or late stage) along with the prediction score (Probability value). The user can use THCA-CN-F for predicting cancer vs normal, THCA-NEL-M for normal vs early vs late, and THCA-All-SVC-L1, THCA-EL-All-WEKA and THCA-EL-PC for predicting early vs late stage.

Another module consists of analysis module which is helpful in evaluating the role of each transcript in discrimination of early stage from the late stage. This module gives p-value (calculated using Wilcoxon rank test) for each transcript that signifies whether the transcript’s expression varies in the early and late stage significantly. It also gives expression threshold and classifying AUROC of each transcript along with average expression of that gene in the early and late stage of cancer. This web server is available from URL “http://webs.iiitd.edu.in/raghava/cancertsp/” for public use.

## Discussion and conclusion

The current study is an attempt to explore the reliable expression-based markers capable of segregating early stage patients from late stage patients. The FNA allows the diagnosis of nature of thyroid nodules in the majority of cases. However, despite the benefits of FNA for diagnosing papillary, medullary, and anaplastic thyroid cancer, it is not helpful in determining whether the thyroid tumors are benign or malignant. In addition, some FNA results suggest, but do not definitively diagnose, papillary thyroid cancer (Grogan et al., 2010). With the advent of genomics technology, publicly available cancer patient’s expression data from resources like TCGA has expedited the search for expression-based molecular markers, capable of reliable diagnosis in clinical settings.

In this work, we made an attempt to understand how well (prediction power in terms of AUROC) the gene expression predicts the stage of the THCA tumor samples. First, we ranked all the transcripts on the basis of AUROC, calculated based on simple expression based threshold. The expression of single gene DCN, at the threshold of 3.01 (log2 FPKM) showed maximum AUROC of 0.66 with mean log2 FPKM of 2.18 in early stage patients and log2 FPKM of 3.03 in late stage patients. This indicates that generally the expression of DCN is less than log2 FPKM of 3.01 in early stage patient. The DCN gene is member of the extracellular small leucine-rich proteoglycan family present in connective tissues. Arnalde et al. showed that DCN can be potential diagnostic marker and therapeutic target for PTC (Arnaldi et al., 2005). It also has been shown that increased expression of DCN leads to decreased adhesion and increased migration of glioma cells by downregulation of TGF-β signaling (Yao et al., 2016). Next, various combinations are tested to elucidate potential biomarkers segregating early and late stage. From 60,483 transcripts we explored various feature spaces like protein coding transcripts, cancer hallmark transcripts and all types of transcripts (protein coding and non-coding transcripts). The SVM model based on THCA-EL-All-WEKA (78 features), selected using WEKA (from coding and non-coding transcripts) resulted in 74.51% accuracy and 0.73 AUROC on validation data. The various types of features in this signature reveal the role of various non-coding transcripts along with protein coding transcripts in progression of cancer. Out of 28 protein coding, five genes; TERT (Liu and Xing, 2016), FLT4 (Tiedje et al., 2016), DUSP6 (Buffet et al., 2017), USP10 (Cui et al., 2014) and POMC (Sheikh-Ali et al., 2007) have already been implicated in thyroid cancer. miR-3196 has also been found to downregulated in PTC non-metastasized patients (Qiu et al., 2015). This shows that many components out of 78 signatures have already been implicated in PTC and other cancers. These genes can be further investigated to reveal their role as biomarkers for early stage PTC. Another model selected using SVC-L1 feature selection resulted in nearly the half the features as compared to 78 features and comparable performance to discriminate early and late stage features. These 36 features lead to similar accuracy and high specificity of 70 % as compared 58.2% using 78 features. One of the most studied TERT gene is part of this signature whose promoter mutations are closely associated with aggressive clinicopathological characteristics and poor prognosis in PTC (Liu et al., 2016). We also validated this signature using external validation data where we were able to predict 70.6% early stage samples and 100% late stage samples using 18 available features from 36 features after applying cross platform normalization. Next, we also applied multiclass machine learning to distinguish normal, early and late samples and could obtain 75.43 % accuracy with 0.80 AUROC on validation data using THCA-NEL-M signature of 211 transcripts. This classification pointed out that mostly late stage samples were misclassified as early stage samples where as normal and early stage sample were classified correctly.

Subsequently, correlation based analysis is performed to identify genes that have disturbed correlation in early stage in comparison to the late stage of cancer. The genes which show disturbed correlation in one stage as compared to other stage mainly belongs to neuroactive ligand receptor interaction pathway.

Next, transcripts having high prediction capability in terms of AUROC to segregate cancer and normal samples also have been derived. Interestingly overlapping sense transcript RP11-363E7.4 showed highest AUROC of 0.96 in classifying cancer samples from normal samples. It has been already shown in literature that sense to antisense transcript ratio increases in Cancer (Maruyama et al., 2012). Other protein coding transcript FAM84A, shows 0.95 AUROC and has already been reported to play role in metastasis of liver and colon cancer (Kobayashi et al., 2006;Kamino et al., 2011). In this study, AUROC of most of the signatures to segregate cancer and normal samples have similar range as reported by earlier studies, which further validates our findings with literature (Cong et al., 2015). Further using five protein coding transcripts (THCA-CL-PC), we were able to classify cancer samples from normal samples with 97.32% accuracy and 0.99 AUROC.

Eventually, a web server CancerTSP is developed where the user can provide transcript’s expression (FPKM values) and can predict whether the cancer is in the early or late stage. This type of application where expression of transcripts is used to demarcate the early and late stage of cancer using machine learning can provide better understandings about the role of diverse transcripts responsible for development of cancer from early to late stage of cancer. Hence, this resource will help the scientific community in making preliminary hypotheses regarding cancer progression.

## Acknowledgement

The authors acknowledge funding agencies J. C. Bose National Fellowship (DST). S.B. and H.K. are thankful to ICMR and CSIR for providing fellowships.

## Authors Contributions

S.B. and H.K. collected the data and processed the datasets. S.B., H.K., R.K., and S.S. developed classification models. S.B. and H.K. created the back-end server and front-end user interface. S.B., H.K. and G.P.S.R. analyzed the results. H.K. and S.B. and G.P.S.R penned the manuscript. G.P.S.R. conceived and coordinated the project, facilitated in the interpretation and data analysis and gave overall supervision to the project. All authors have read and approved the final manuscript.

## References

Antal, C.E., Hudson, A.M., Kang, E., Zanca, C., Wirth, C., Stephenson, N.L., Trotter, E.W., Gallegos, L.L., Miller, C.J., Furnari, F.B., Hunter, T., Brognard, J., and Newton, A.C. (2015). Cancer-associated protein kinase C mutations reveal kinase’s role as tumor suppressor. Cell 160, 489–502.

Arnaldi, L.A., Borra, R.C., Maciel, R.M., and Cerutti, J.M. (2005). Gene expression profiles reveal that DCN, DIO1, and DIO2 are underexpressed in benign and malignant thyroid tumors. Thyroid 15, 210–221.

Baraniuk, R.G. (2007). Compressive Sensing [Lecture Notes]. IEEE Signal Processing Magazine 24, 118 – 121.

Basolo, F., Fugazzola, L., Fontanini, G., Elisei, R., Pepe, S., Bevilacqua, G., Pinchera, A., and Pacini, F. (1993). Markers of cell-proliferation as prognostic factors in differentiated thyroid-cancer. Int J Oncol 3, 1077–1081.

Bhalla, S., Chaudhary, K., Kumar, R., Sehgal, M., Kaur, H., Sharma, S., and Raghava, G.P. (2017). Gene expression-based biomarkers for discriminating early and late stage of clear cell renal cancer. Sci Rep 7, 44997.

Bhondekar, A., Kaur, R., Kumar, R., Vig, R., and Kapur, P. (2011). A novel approach using Dynamic Social Impact Theory for optimization of impedance-Tongue (iTongue). Chemometrics and Intelligent Laboratory Systems 109, 65–76.

Borrelli, N., Ugolini, C., Giannini, R., Antonelli, A., Giordano, M., Sensi, E., Torregrossa, L., Fallahi, P., Miccoli, P., and Basolo, F. (2016). Role of gene expression profiling in defining indeterminate thyroid nodules in addition to BRAF analysis. Cancer Cytopathol 124, 340–349.

Buffet, C., Hecale-Perlemoine, K., Bricaire, L., Dumont, F., Baudry, C., Tissier, F., Bertherat, J., Cochand-Priollet, B., Raffin-Sanson, M.L., Cormier, F., and Groussin, L. (2017). DUSP5 and DUSP6, two ERK specific phosphatases, are markers of a higher MAPK signaling activation in BRAF mutated thyroid cancers. PloS One 12, e0184861.

Cancer Genome Atlas Research, N. (2014). Integrated genomic characterization of papillary thyroid carcinoma. Cell 159, 676–690.

Chai, Y.J., Yi, J.W., Jee, H.G., Kim, Y.A., Kim, J.H., Xing, M., and Lee, K.E. (2016). Significance of the BRAF mRNA Expression Level in Papillary Thyroid Carcinoma: An Analysis of The Cancer Genome Atlas Data. PloS One 11, e0159235.

Chedotal, A. (2007). Chemotropic axon guidance molecules in tumorigenesis. Prog Exp Tumor Res 39, 78–90.

Chen, E.Y., Tan, C.M., Kou, Y., Duan, Q., Wang, Z., Meirelles, G.V., Clark, N.R., and Ma’ayan, A. (2013). Enrichr: interactive and collaborative HTML5 gene list enrichment analysis tool. BMC Bioinformatics 14, 128.

Chiu, C.G., Strugnell, S.S., Griffith, O.L., Jones, S.J., Gown, A.M., Walker, B., Nabi, I.R., and Wiseman, S.M. (2010). Diagnostic utility of galectin-3 in thyroid cancer. Am J Pathol 176, 2067–2081.

Choi, J.Y., Yi, J.W., Lee, J.H., Song, R.Y., Yu, H., Kwon, H., Chai, Y.J., Kim, S.J., and Lee, K.E. (2017). VDR mRNA overexpression is associated with worse prognostic factors in papillary thyroid carcinoma. Endocr Connect 6, 172–178.

Chudova, D., Wilde, J.I., Wang, E.T., Wang, H., Rabbee, N., Egidio, C.M., Reynolds, J., Tom, E., Pagan, M., Rigl, C.T., Friedman, L., Wang, C.C., Lanman, R.B., Zeiger, M., Kebebew, E., Rosai, J., Fellegara, G., Livolsi, V.A., and Kennedy, G.C. (2010). Molecular classification of thyroid nodules using high-dimensionality genomic data. J Clin Endocrinol Metab 95, 5296–5304.

Clough, E., and Barrett, T. (2016). The Gene Expression Omnibus Database. Methods Mol Biol 1418, 93–110.

Cong, D., He, M., Chen, S., Liu, X., Liu, X., and Sun, H. (2015). Expression profiles of pivotal microRNAs and targets in thyroid papillary carcinoma: an analysis of The Cancer Genome Atlas. Onco Targets Ther 8, 2271–2277.

Cui, B., Yang, Q., Guan, H., Shi, B., Hou, P., and Ji, M. (2014). PRIMA-1, a mutant p53 reactivator, restores the sensitivity of TP53 mutant-type thyroid cancer cells to the histone methylation inhibitor 3-Deazaneplanocin A. J Clin Endocrinol Metab 99, E962–970.

Cui, H., Zhang, Y., Zhang, Q., Chen, W., Zhao, H., and Liang, J. (2017). A comprehensive genome-wide analysis of long noncoding RNA expression profile in hepatocellular carcinoma. Cancer Med 6, 2932–2941.

De Groot, J.F., Piao, Y., Lu, L., Fuller, G.N., and Yung, W.K. (2008). Knockdown of GluR1 expression by RNA interference inhibits glioma proliferation. J Neurooncol 88, 121–133.

Dohi, O., Takada, H., Wakabayashi, N., Yasui, K., Sakakura, C., Mitsufuji, S., Naito, Y., Taniwaki, M., and Yoshikawa, T. (2010). Epigenetic silencing of RELN in gastric cancer. Int J Oncol 36, 85–92.

Eibe Frank, M.a.H., and Ian H. Witten (2016). The WEKA Workbench. Online Appendix for “Data Mining: Practical Machine Learning Tools and Techniques. Morgan Kaufmann.

Fan, M., Li, X., Jiang, W., Huang, Y., Li, J., and Wang, Z. (2013). A long non-coding RNA, PTCSC3, as a tumor suppressor and a target of miRNAs in thyroid cancer cells. Exp Ther Med 5, 1143–1146.

Frank, E., Hall, M., Trigg, L., Holmes, G., and Witten, I.H. (2004). Data mining in bioinformatics using Weka. Bioinformatics 20, 2479–2481.

Franks, J.M., Cai, G., and Whitfield, M.L. (2018). Feature specific quantile normalization enables cross-platform classification of molecular subtypes using gene expression data. Bioinformatics 34, 1868–1874.

Fresno Vara, J.A., Casado, E., De Castro, J., Cejas, P., Belda-Iniesta, C., and Gonzalez-Baron, M. (2004). PI3K/Akt signalling pathway and cancer. Cancer Treat Rev 30, 193–204.

Gharib, H. (1994). Fine-needle aspiration biopsy of thyroid nodules: advantages, limitations, and effect. Mayo Clin Proc 69, 44–49.

Grogan, R.H., Mitmaker, E.J., and Clark, O.H. (2010). The evolution of biomarkers in thyroid cancer-from mass screening to a personalized biosignature. Cancers (Basel) 2, 885–912.

Haider, A.S., Rakha, E.A., Dunkley, C., and Zaitoun, A.M. (2011). The impact of using defined criteria for adequacy of fine needle aspiration cytology of the thyroid in routine practice. Diagn Cytopathol 39, 81–86.

Hanahan, D., and Weinberg, R.A. (2011). Hallmarks of cancer: the next generation. Cell 144, 646–674.

Hay, I.D., Thompson, G.B., Grant, C.S., Bergstralh, E.J., Dvorak, C.E., Gorman, C.A., Maurer, M.S., Mciver, B., Mullan, B.P., Oberg, A.L., Powell, C.C., Van Heerden, J.A., and Goellner, J.R. (2002). Papillary thyroid carcinoma managed at the Mayo Clinic during six decades (1940-1999): temporal trends in initial therapy and long-term outcome in 2444 consecutively treated patients. World J Surg 26, 879–885.

Herner, A., Sauliunaite, D., Michalski, C.W., Erkan, M., De Oliveira, T., Abiatari, I., Kong, B., Esposito, I., Friess, H., and Kleeff, J. (2011). Glutamate increases pancreatic cancer cell invasion and migration via AMPA receptor activation and Kras-MAPK signaling. Int J Cancer 129, 2349–2359.

Hwang, B.J., Jin, J., Gunther, R., Madabushi, A., Shi, G., Wilson, G.M., and Lu, A.L. (2015). Association of the Rad9-Rad1-Hus1 checkpoint clamp with MYH DNA glycosylase and DNA. DNA Repair (Amst) 31, 80–90.

Ishitani, T., Kishida, S., Hyodo-Miura, J., Ueno, N., Yasuda, J., Waterman, M., Shibuya, H., Moon, R.T., Ninomiya-Tsuji, J., and Matsumoto, K. (2003). The TAK1-NLK mitogen-activated protein kinase cascade functions in the Wnt-5a/Ca(2+) pathway to antagonize Wnt/beta-catenin signaling. Mol Cell Biol 23, 131–139.

Kamino, H., Yamazaki, Y., Saito, K., Takizawa, D., Kakizaki, S., Moore, R., and Negishi, M. (2011). Nuclear receptor CAR-regulated expression of the FAM84A gene during the development of mouse liver tumors. Int J Oncol 38, 1511–1520.

Kaur, R., Kumar R, Bhondekar, A., and Kapur, P. (2013). Human opinion dynamics: An inspiration to solve complex optimization problems. Scientific Reports 3, 3008.

Kaur, R., Kumar, R., Gulati, A., Ghanshyam, C., Kapur, P., and Bhondekar, A. (2012). Enhancing electronic nose performance: A novel feature selection approach using dynamic social impact theory and moving window time slicing for classification of Kangra orthodox black tea (Camellia sinensis (L.) O. Kuntze). Sensors Actuators B Chem 166, 309–319.

Kim, S.J., Park, J.W., Yoon, J.S., Mok, J.O., Kim, Y.J., Park, H.K., Kim, C.H., Byun, D.W., Lee, Y.J., Jin, S.Y., Suh, K.I., and Yoo, M.H. (2004). Increased expression of focal adhesion kinase in thyroid cancer: immunohistochemical study. J Korean Med Sci 19, 710–715.

Kobayashi, T., Masaki, T., Sugiyama, M., Atomi, Y., Furukawa, Y., and Nakamura, Y. (2006). A gene encoding a family with sequence similarity 84, member A (FAM84A) enhanced migration of human colon cancer cells. Int J Oncol 29, 341–347.

Korc, M. (2010). Driver mutations: a roadmap for getting close and personal in pancreatic cancer. Cancer Biol Ther 10, 588–591.

Kuleshov, M.V., Jones, M.R., Rouillard, A.D., Fernandez, N.F., Duan, Q., Wang, Z., Koplev, S., Jenkins, S.L., Jagodnik, K.M., Lachmann, A., Mcdermott, M.G., Monteiro, C.D., Gundersen, G.W., and Ma’ayan, A. (2016). Enrichr: a comprehensive gene set enrichment analysis web server 2016 update. Nucleic Acids Res 44, W90–97.

Liberzon, A., Birger, C., Thorvaldsdottir, H., Ghandi, M., Mesirov, J.P., and Tamayo, P. (2015). The Molecular Signatures Database (MSigDB) hallmark gene set collection. Cell Syst 1, 417–425.

Liu, C., Liu, Z., Chen, T., Zeng, W., Guo, Y., and Huang, T. (2016). TERT promoter Mutation and Its Association with Clinicopathological Features and Prognosis of Papillary Thyroid Cancer: A Meta-analysis. Sci Rep 6, 36990.

Liu, R., and Xing, M. (2016). TERT promoter mutations in thyroid cancer. Endocr Relat Cancer 23, R143–155.

Liu, R., Zhang, T., Zhu, G., and Xing, M. (2018). Regulation of mutant TERT by BRAF V600E/MAP kinase pathway through FOS/GABP in human cancer. Nat Commun 9, 579.

Liu, T.J., Koul, D., Lafortune, T., Tiao, N., Shen, R.J., Maira, S.M., Garcia-Echevrria, C., and Yung, W.K. (2009). NVP-BEZ235, a novel dual phosphatidylinositol 3-kinase/mammalian target of rapamycin inhibitor, elicits multifaceted antitumor activities in human gliomas. Mol Cancer Ther 8, 2204–2210.

Martins, M.B., Marcello, M.A., Batista, F.A., Peres, K.C., Meneghetti, M., Ward, M.a.L., Etchebehere, E., Da Assumpcao, L.V.M., and Ward, L.S. (2018). Serum interleukin measurement may help identify thyroid cancer patients with active disease. Clin Biochem 52, 1–7.

Maruyama, R., Shipitsin, M., Choudhury, S., Wu, Z., Protopopov, A., Yao, J., Lo, P.K., Bessarabova, M., Ishkin, A., Nikolsky, Y., Liu, X.S., Sukumar, S., and Polyak, K. (2012). Altered antisense-to-sense transcript ratios in breast cancer. Proc Natl Acad Sci U S A 109, 2820–2824.

Matson, D.R., Hardin, H., Buehler, D., and Lloyd, R.V. (2017). AKT activity is elevated in aggressive thyroid neoplasms where it promotes proliferation and invasion. Exp Mol Pathol 103, 288–293.

Mazzoni, E., Adam, A., Bal De Kier Joffe, E., and Aguirre-Ghiso, J.A. (2003). Immortalized mammary epithelial cells overexpressing protein kinase C gamma acquire a malignant phenotype and become tumorigenic in vivo. Mol Cancer Res 1, 776–787.

Mohammed, H., Russell, I.A., Stark, R., Rueda, O.M., Hickey, T.E., Tarulli, G.A., Serandour, A.A., Birrell, S.N., Bruna, A., Saadi, A., Menon, S., Hadfield, J., Pugh, M., Raj, G.V., Brown, G.D., D’santos, C., Robinson, J.L., Silva, G., Launchbury, R., Perou, C.M., Stingl, J., Caldas, C., Tilley, W.D., and Carroll, J.S. (2015). Corrigendum: Progesterone receptor modulates ERalpha action in breast cancer. Nature 526, 144.

Morris, L.G., Shaha, A.R., Tuttle, R.M., Sikora, A.G., and Ganly, I. (2010). Tall-cell variant of papillary thyroid carcinoma: a matched-pair analysis of survival. Thyroid 20, 153–158.

Myatt, S.S., and Lam, E.W. (2007). The emerging roles of forkhead box (Fox) proteins in cancer. Nat Rev Cancer 7, 847–859.

Nga, M.E., Lim, G.S., Soh, C.H., and Kumarasinghe, M.P. (2008). HBME-1 and CK19 are highly discriminatory in the cytological diagnosis of papillary thyroid carcinoma. Diagn Cytopathol 36, 550–556.

Niu, X.M., and Lu, S. (2014). Acetylcholine receptor pathway in lung cancer: New twists to an old story. World J Clin Oncol 5, 667–676.

Okamura, Y., Nomoto, S., Kanda, M., Hayashi, M., Nishikawa, Y., Fujii, T., Sugimoto, H., Takeda, S., and Nakao, A. (2011). Reduced expression of reelin (RELN) gene is associated with high recurrence rate of hepatocellular carcinoma. Ann Surg Oncol 18, 572–579.

Parsons, M., and Adams, J.C. (2008). Rac regulates the interaction of fascin with protein kinase C in cell migration. J Cell Sci 121, 2805–2813.

Pavlopoulos, G.A., Secrier, M., Moschopoulos, C.N., Soldatos, T.G., Kossida, S., Aerts, J., Schneider, R., and Bagos, P.G. (2011). Using graph theory to analyze biological networks. BioData Min 4, 10.

Pedregosa, F.a.V. G. and Gramfort, A. and Michel, V., and Thirion, B.a.G. O. and Blondel, M. and Prettenhofer, P., and Weiss, R.a.D. V. and Vanderplas, J. and Passos, A. And, and Cournapeau, D.a.B. M. and Perrot, M. and Duchesnay, E. (2011). Scikit-learn: Machine Learning in Python. Journal of Machine Learning Research 12, 2825-2830.

Pellegriti, G., Frasca, F., Regalbuto, C., Squatrito, S., and Vigneri, R. (2013). Worldwide increasing incidence of thyroid cancer: update on epidemiology and risk factors. J Cancer Epidemiol 2013, 965212.

Qiu, Z.L., Shen, C.T., Song, H.J., Wei, W.J., and Luo, Q.Y. (2015). Differential expression profiling of circulation microRNAs in PTC patients with non-131I and 131I-avid lungs metastases: a pilot study. Nucl Med Biol 42, 499–504.

Salomaki, H.H., Sainio, A.O., Soderstrom, M., Pakkanen, S., Laine, J., and Jarvelainen, H.T. (2008). Differential expression of decorin by human malignant and benign vascular tumors. J Histochem Cytochem 56, 639–646.

Savari, S., Liu, M., Zhang, Y., Sime, W., and Sjolander, A. (2013). CysLT(1)R antagonists inhibit tumor growth in a xenograft model of colon cancer. PloS One 8, e73466.

Sethi, K., Sarkar, S., Das, S., Mohanty, B., and Mandal, M. (2010). Biomarkers for the diagnosis of thyroid cancer. J Exp Ther Oncol 8, 341–352.

Shannon, P., Markiel, A., Ozier, O., Baliga, N.S., Wang, J.T., Ramage, D., Amin, N., Schwikowski, B., and Ideker, T. (2003). Cytoscape: a software environment for integrated models of biomolecular interaction networks. Genome Res 13, 2498–2504.

Sheikh-Ali, M., Krishna, M., Lloyd, R., and Smallridge, R.C. (2007). Predicting the development of Cushing’s syndrome in medullary thyroid cancer: utility of proopiomelanocortin messenger ribonucleic acid in situ hybridization. Thyroid 17, 631–634.

Smallridge, R.C., Chindris, A.M., Asmann, Y.W., Casler, J.D., Serie, D.J., Reddi, H.V., Cradic, K.W., Rivera, M., Grebe, S.K., Necela, B.M., Eberhardt, N.L., Carr, J.M., Mciver, B., Copland, J.A., and Thompson, E.A. (2014). RNA sequencing identifies multiple fusion transcripts, differentially expressed genes, and reduced expression of immune function genes in BRAF (V600E) mutant vs BRAF wild-type papillary thyroid carcinoma. J Clin Endocrinol Metab 99, E338–347.

Song, Y., Li, L., Ou, Y., Gao, Z., Li, E., Li, X., Zhang, W., Wang, J., Xu, L., Zhou, Y., Ma, X., Liu, L., Zhao, Z., Huang, X., Fan, J., Dong, L., Chen, G., Ma, L., Yang, J., Chen, L., He, M., Li, M., Zhuang, X., Huang, K., Qiu, K., Yin, G., Guo, G., Feng, Q., Chen, P., Wu, Z., Wu, J., Ma, L., Zhao, J., Luo, L., Fu, M., Xu, B., Chen, B., Li, Y., Tong, T., Wang, M., Liu, Z., Lin, D., Zhang, X., Yang, H., Wang, J., and Zhan, Q. (2014). Identification of genomic alterations in oesophageal squamous cell cancer. Nature 509, 91–95.

Stephen, J.K., Chen, K.M., Merritt, J., Chitale, D., Divine, G., and Worsham, M.J. (2015). Methylation Markers for Early Detection and Differentiation of Follicular Thyroid Cancer Subtypes. Cancer Clin Oncol 4, 1–12.

Stokowy, T., Gawel, D., and Wojtas, B. (2016). Differences in miRNA and mRNA Profile of Papillary Thyroid Cancer Variants. Int J Endocrinol 2016, 1427042.

Szklarczyk, D., Franceschini, A., Wyder, S., Forslund, K., Heller, D., Huerta-Cepas, J., Simonovic, M., Roth, A., Santos, A., Tsafou, K.P., Kuhn, M., Bork, P., Jensen, L.J., and Von Mering, C. (2015). STRING v10: protein-protein interaction networks, integrated over the tree of life. Nucleic Acids Res 43, D447–452.

Theocharis, A.D., and Karamanos, N.K. (2017). Proteoglycans remodeling in cancer: Underlying molecular mechanisms. Matrix Biol.

Tiedje, V., Ting, S., Walter, R.F., Herold, T., Worm, K., Badziong, J., Zwanziger, D., Schmid, K.W., and Fuhrer, D. (2016). Prognostic markers and response to vandetanib therapy in sporadic medullary thyroid cancer patients. Eur J Endocrinol 175, 173–180.

Tomczak, K., Czerwinska, P., and Wiznerowicz, M. (2015). The Cancer Genome Atlas (TCGA): an immeasurable source of knowledge. Contemp Oncol (Pozn) 19, A68–77.

Von Bergh, A.R., Van Drunen, E., Van Wering, E.R., Van Zutven, L.J., Hainmann, I., Lonnerholm, G., Meijerink, J.P., Pieters, R., and Beverloo, H.B. (2006). High incidence of t(7;12)(q36;p13) in infant AML but not in infant ALL, with a dismal outcome and ectopic expression of HLXB9. Genes Chromosomes Cancer 45, 731–739.

Wilkens, L., Jaggi, R., Hammer, C., Inderbitzin, D., Giger, O., and Von Neuhoff, N. (2011). The homeobox gene HLXB9 is upregulated in a morphological subset of poorly differentiated hepatocellular carcinoma. Virchows Arch 458, 697– 708.

Xu, C., Qi, X., Du, X., Zou, H., Gao, F., Feng, T., Lu, H., Li, S., An, X., Zhang, L., Wu, Y., Liu, Y., Li, N., Capecchi, M.R., and Wu, S. (2017). piggyBac mediates efficient in vivo CRISPR library screening for tumorigenesis in mice. Proc Natl Acad Sci U S A 114, 722–727.

Yao, T., Zhang, C.G., Gong, M.T., Zhang, M., Wang, L., and Ding, W. (2016). Decorin-mediated inhibition of the migration of U87MG glioma cells involves activation of autophagy and suppression of TGF-beta signaling. FEBS Open Bio 6, 707–719.

Yi, J.W., Kim, S.J., Kim, J.K., Seong, C.Y., Yu, H.W., Chai, Y.J., Choi, J.Y., and Lee, K.E. (2017). Upregulation of the ESR1 Gene and ESR Ratio (ESR1/ESR2) is Associated with a Worse Prognosis in Papillary Thyroid Carcinoma: The Impact of the Estrogen Receptor alpha/beta Expression on Clinical Outcomes in Papillary Thyroid Carcinoma Patients. Ann Surg Oncol 24, 3754–3762.

Yu, H., Xu, Q., Liu, F., Ye, X., Wang, J., and Meng, X. (2015). Identification and validation of long noncoding RNA biomarkers in human non-small-cell lung carcinomas. J Thorac Oncol 10, 645–654.

Yu L L.H. (2003). Feature selection for high-dimensional data: A fast correlation-based filter solution. ICML, 856–863.

Zhang, L., Wang, J., Wang, Y., Zhang, Y., Castro, P., Shao, L., Sreekumar, A., Putluri, N., Guha, N., Deepak, S., Padmanaban, A., Creighton, C.J., and Ittmann, M. (2016). MNX1 Is Oncogenically Upregulated in African-American Prostate Cancer. Cancer Res 76, 6290–6298.

Zolotov, S. (2016). Genetic Testing in Differentiated Thyroid Carcinoma: Indications and Clinical Implications. Rambam Maimonides Med J 7.

